# Oxytocin increases trust in humans with a low disposition to trust

**DOI:** 10.1101/2025.10.01.679711

**Authors:** Bodo Vogt, Paul Bengart, Caroylyn Declerck, Ernst Fehr

## Abstract

In recent years, increasing skepticism regarding oxytocin’s (OT) influence on social behavior arose. Low power, HARKing (hypothesizing after the results are known), and replication failures have clouded the field. Here, we directly address these concerns with a high-powered, preregistered study that offers robust evidence for a causal effect of OT on trust among individuals with a low disposition to trust. We recruited 359 low-trusting individuals who participated in a trust game under strict anonymity conditions. Results show that OT administration significantly increased trusting behavior by roughly 15%, with consistent effects across regression models with and without controls for personality traits. A pooled data analysis incorporating a previous sample (n=219) of low-trusting individuals further strengthens this conclusion, yielding a statistically significant 16.9% increase in trust. Crucially, no interaction effect was found between OT and the degree of dispositional trust, suggesting OT’s effect is uniform across the low-trusting spectrum. These findings present a strong case for OT’s selective trust-enhancing role. By isolating OT’s impact within a well-defined subpopulation and experimental context, this study provides a critical pivot in the debate over neurobiological mechanisms of trust.

---

Twenty years after the first publication suggesting that the neuropeptide oxytocin (OT) increases trust in humans^1^, the debate regarding the validity of this statement remains unresolved. Yet the question continues to be of significant interest because trust is the foundation of individual well-being, healthy social relationships, economic exchanges, and effective governance. A deeper understanding of the biological underpinnings of trust may therefore help to sustain these important pillars of society.

Several factors have cast doubt on the proposed causal link between oxytocin (OT) and trust, as well as on OT’s broader role in regulating human social behavior. First, The abundance of underpowered studies examining intranasal OT’s effects on human psychology and social behaviors such as trust, empathy, emotion recognition, and mind reading, has led to numerous null findings, which may have contributed to a “file drawer problem”^2^, meaning that the current state of the art might be biased because null findings typically are not published.

Second, when initial hypotheses fail to be supported, exploratory analyses often reveal interaction effects. However, these post-hoc findings still lack the statistical power to be conclusive. Moreover, they are rarely replicated. Mierop et al.^3^ estimate that up to 90% of reported OT interaction effects may be false positives, while the intended main effects remain undetectable due to small sample sizes. None of the studies with a between-subject design in the review by Mierop et al.^3^ reached a power of 50%.

Third, meta-analyses, mostly focused on clinical populations, yielded mixed results^4–6^. The only meta-analysis on intranasal OT and trust found no effect^7^, though it included studies with questionable or highly varied methodologies that cannot be considered true replications^8^.

As a result, a growing number of critical reviews have questioned the usefulness of intranasal OT in understanding human psychology and social behavior and proposed methodological improvements^2,3,9–17^. Today, these critical reviews even seem to outnumber genuine replication attempts. A 10-year literature search in the Web of Science using "intranasal oxytocin" and "replication" found only a handful of replication studies specifically addressing the causal impact of OT on human psychology and social behavior^8,18–21^.

Against this backdrop, Declerck et al.^8^ published the results of the first large-scale (N=677), pre-registered replication study re-examining the causal influence of intranasal OT on young adult men’s trusting behavior in the trust game ^1^. The 2020 experiment was conducted in two conditions, and in each of the conditions, the power to detect an effect size of OT like the one observed in Kosfeld et al.^1^ was 99% - based on a one-sided t-test with a significance threshold of α = 0.05. The first condition (labelled “social contact condition”) replicated as close as possible the original Kosfeld et al.^1^ study, in which participants briefly intermingled while seated at communal tables before participating in a trust game with an unknown counterpart from the group. It was hypothesized that in this condition minimal social cues could create a context conducive to bonding and trust-building (in line with the social salience hypothesis of OT^22^). In the second condition, participants were seated in cubicles throughout the experiment, ensuring complete anonymity (“no-contact condition”) of the interaction partners in the trust game. Another important characteristic of the study concerned the assessment of an independent measure of participants’ baseline propensity to trust that took place approximately one week before subjects participated in the trust game. This measure enables the separate examination of OT’s effects on trust for those with a low and those with a high disposition to trust.

Three key findings emerged from Declerck et al.^8^, offering important insights for future research: First, in both conditions no overall effect of OT was observed when considering the entire population. Second, although Declerck et al.^8^ conjectured – based on the findings and the design of the Kosfeld et al.^1^ study – that OT is more likely to have a trust-enhancing effect in the social contact condition, they did not find such an effect. In contrast, they observed a negative interaction between OT and the social contact condition that is particularly large and significant for the subpopulation of individuals with a low propensity to trust. While OT may have enhanced the positive valence of social cues and facilitated trust in the original Kosfeld et al.^1^ study, the results of the 2020 study inadvertently may have offset this effect, as participants generally reported feeling more disconnected from each other in the social contact condition which is the opposite of a trust-facilitating environment. OT is presumably not likely to facilitate trusting behaviors in such an environment^22^.

Third, in the no-contact condition, *exploratory analyses* revealed a significant effect of OT in a subpopulation of participants (N=191) with a low disposition to trust. No such effect was observed in participants with a high disposition to trust, which may have been due to a ceiling effect. The high trustors typically invested already 80 % of their endowment in the placebo condition, leaving little room to detect additional treatment effects. This result parallels findings in pharmacological studies where, for example, a certain medicinal drug may have an analgesic effect, but only in patients who are truly experiencing pain. The drug brings no relief to healthy people who are not suffering.

However, while finding a significant effect of OT for low-trusting individuals in the no-contact condition is interesting and encouraging, two aspects of this result may dampen an observer’s enthusiasm. First, the finding resulted from a post-hoc exploratory analysis and was not part of the pre-registered set of hypotheses. Second, the finding is based on a statistical power of 60% for a two-sided test and a significance level of α = 0.05, which is considerably above the typical power of OT administration studies reported by Walum et al. ^15^ but still not overwhelming.

In the current study we intend to address these reasons for skepticism and examine the empirical validity and robustness of the findings with an unbiased new study that is based on a considerably higher statistical power and a transparently communicated ex-ante hypothesis. Using the initial design and analysis in Declerck et al.^8^ as script for a preregistered study, we thus respond to the growing concerns about HARKing (hypothesizing after the results are known)^23,24^. We preregistered the hypothesis that OT increases trusting behavior of individuals with a low disposition to trust in the no-contact condition on the Open Science Framework (see https://osf.io/bks46/resources), with an intended sample size of n=358 low-trusting individuals (see methods section for sample size calculation). With this sample size, the power to detect an effect size of OT on trust of d = 0.348, which corresponds to the one observed in the no-contact condition of Declerck et al.^8^, is 95% based on a one-sided p-value of P ≤ 0.05.

There were two parts in the current study: a recruitment phase and a laboratory experiment. To recruit participants with a low disposition to trust, we relied on the generalized trust scale^25^ that was embedded in an online questionnaire that had to be filled out approximately one week before the trust experiment. Only individuals who scored 38 or below on this scale – which corresponds to the trust disposition level of all subjects that are at or below the median disposition in Declerck et al.^8^ – were invited to participate in the laboratory trust game.

Those invited to the trust experiment were then randomly assigned to a placebo or OT condition based on a double-blind between-subjects design. We implemented the same trust experiment as in the no-contact condition of Declerck et al.^8^. All participants were assigned the role of investor and informed that they would be randomly matched to a trustee (see methods section). Both investor and trustee received an initial endowment of 12 euros. The investor could choose to send 0, 4, 8, or 12 € to the trustee, knowing that the amount sent would be tripled by the experimenter before adding it to the 12 € endowment of the trustee. The trustee could then reciprocate by returning any amount to the investor. The back-transfer was not tripled. The greater the sum entrusted by the investor, the higher the scope for back-transfers. However, when a trustee is untrustworthy and returns little or nothing, the investor would be worse off than if he had not entrusted any money at all. Thus, the investor’s transfer level is a behavioral indicator of the investor’s trust in receiving a fair back-transfer.

Although it was cumbersome and time consuming to attract a large number of individuals with a low disposition to trust – who are, almost by definition, not easily recruitable – we managed to conduct the trust experiment with n = 359 individuals. In the results section below we first examine the behavior of this sample.

In addition, to further increase the statistical power, we conducted a second set of analyses, where we combine the results of the current experiment with the 219 participants of the no-contact condition of the Declerck et al.^8^ study, who had a dispositional trust score ≤ 38, which gives us n=578. This yields a power of 94.5% to detect an effect of OT on trust like the one in the no-contact condition of Declerck et al.^8^ (i.e., d = 0.348) even with a two-sided p-value of P ≤ 0.05.

## RESULTS from the current study (n=359)

The preregistered hypothesis (see https://osf.io/bks46/resources) that OT increases the transfers of low-trusting investors in the trust game is corroborated by the data. OT increases investors’ trust by 15.1%. Specifically, investors entrusted an average amount of 8.29 (s.d.= 3.97) to the trustees after OT administration, while they only transferred an amount of 7.20 (s.d.= 4.33) in the placebo condition (see Figure 1a; two-sample t-test, two-tailed, t = -2.49, P = 0.0066). We also note that these results are very similar to those reported for the low-trustors by Declerck et al.^8^ in the no-contact condition where the investors transferred 7.10 in the placebo treatment and 8.45 in the OT treatment. Looking at the distribution of investments (see Figure 1b), we also see that, compared to placebo condition, OT administration increases the proportions of €12 and €8 investments while decreasing €4 and €0 investments.

**Figure 1.**
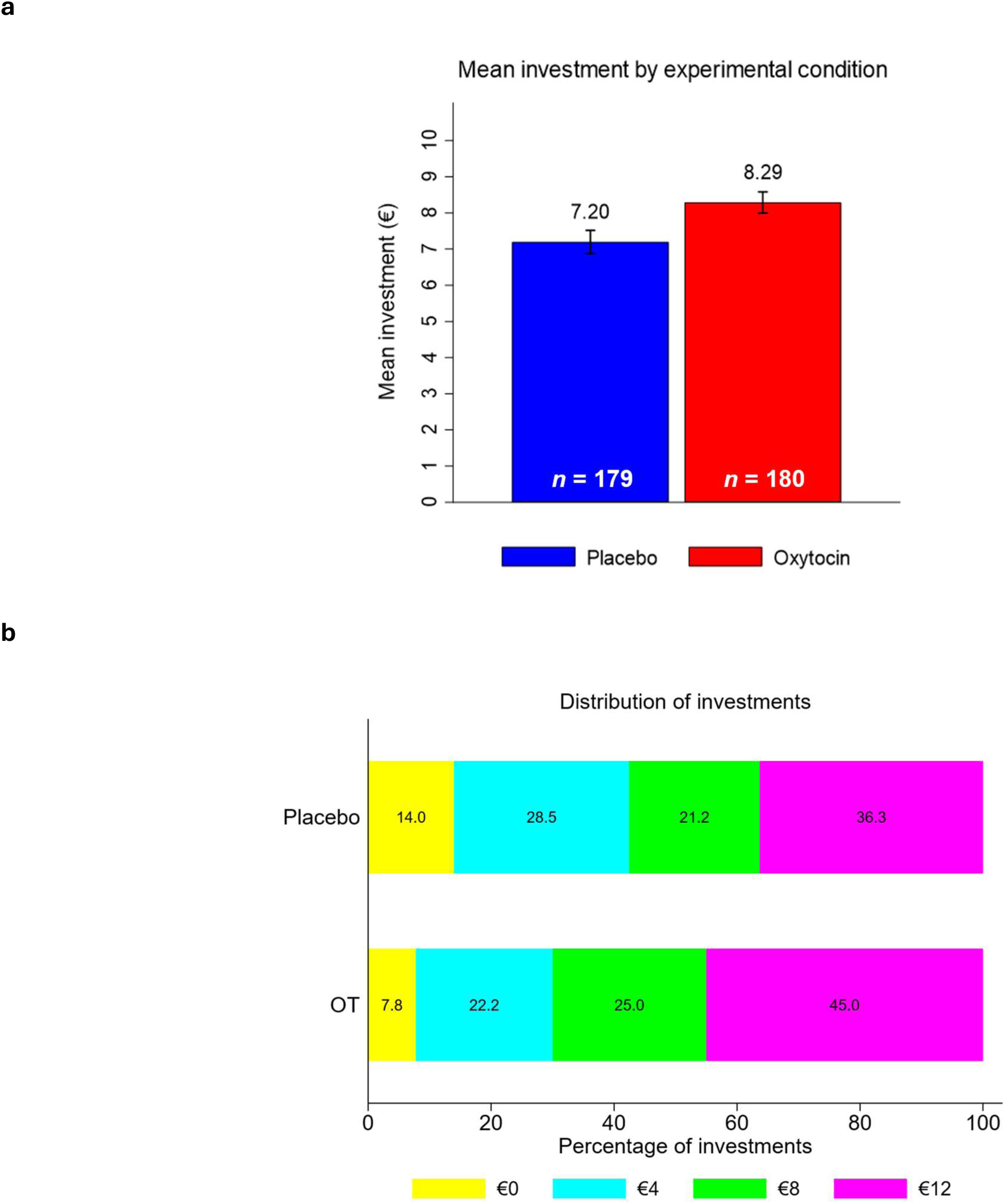
Investment behavior in the trust game: placebo (n = 179) vs OT group (n = 180). **a,** Mean investments in € reported above each bar. Error bars represent standard errors. **b,** Relative frequencies of investing € 0, 4, 8 or 12. Values within the horizontal bars are the percentages of participants who invested these amounts.

To further substantiate these results, we conducted linear regressions with the investors’ transfers as the dependent variable, and a dummy variable for the OT-treatment that captures the effect of OT on transfers as the main independent variable (see Table 1, model 0). In addition, we include the individual’s disposition to trust (model 1) and willingness to take risks (model 2) as covariates, both of which were measured in an online questionnaire approximately one week before the experiment. In model 3 we also controlled for personality variables based on the HEXACO personality inventory^26^. We included the personality factors honesty-humility, the anxiety dimension of emotionality (both of which were measured in the online questionnaire), and extraversion, agreeableness, conscientiousness and openness (measured on the day of the experiment) into the regression analyses. Finally, we also included the extent to which participants felt connected to their interaction partners in the trust game.

**Table 1.**
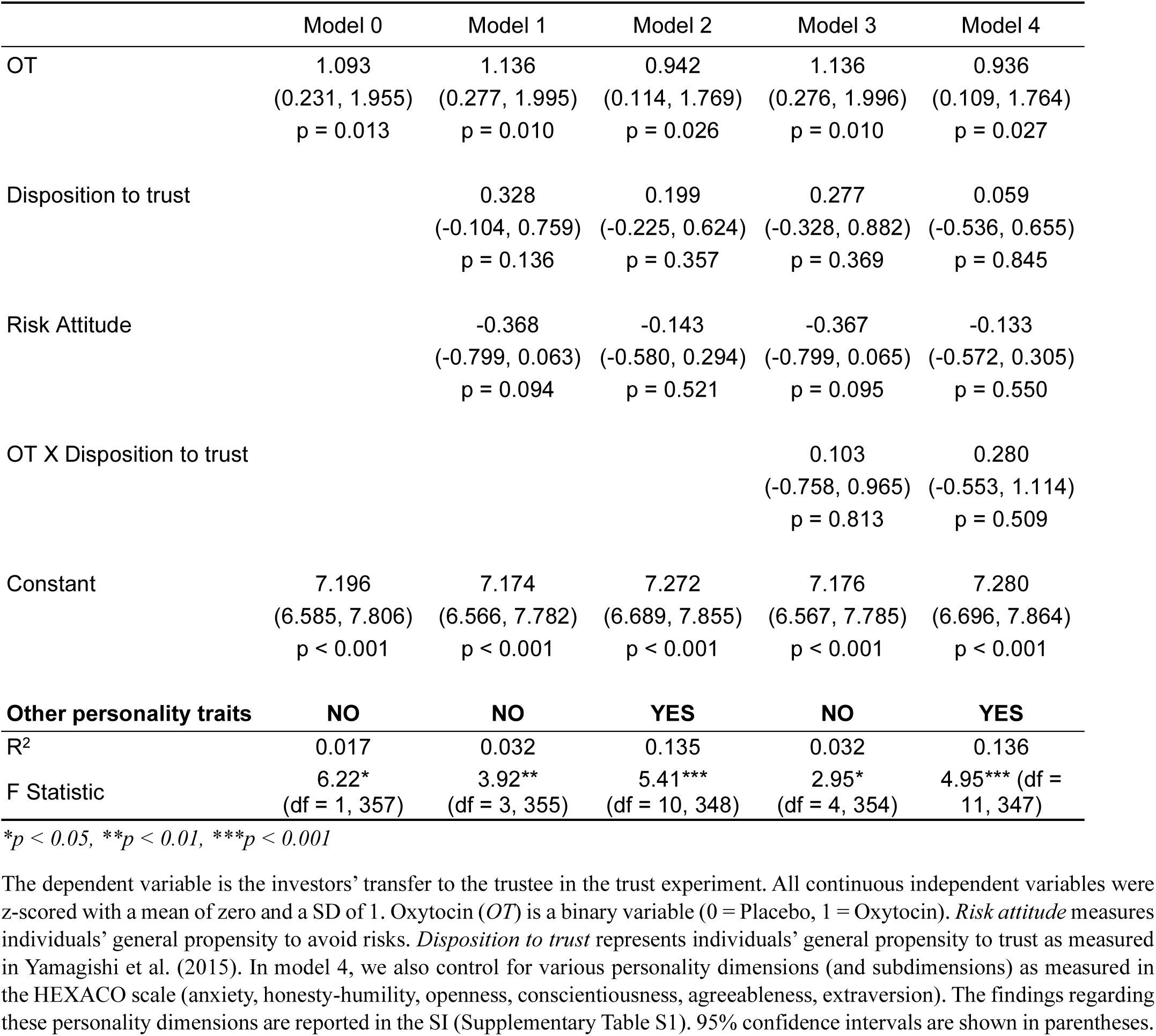
The impact of oxytocin (OT) on investors’ transfers when controlling for covariates in the current study (n = 359)

|  | Model 0 | Model 1 | Model 2 | Model 3 | Model 4 |
| --- | --- | --- | --- | --- | --- |
| OT | 1.093<br>(0.231, 1.955)<br>p = 0.013 | 1.136<br>(0.277, 1.995)<br>p = 0.010 | 0.942<br>(0.114, 1.769)<br>p = 0.026 | 1.136<br>(0.276, 1.996)<br>p = 0.010 | 0.936<br>(0.109, 1.764)<br>p = 0.027 |
| Disposition to trust |  | 0.328<br>(-0.104, 0.759)<br>p = 0.136 | 0.199<br>(-0.225, 0.624)<br>p = 0.357 | 0.277<br>(-0.328, 0.882)<br>p = 0.369 | 0.059<br>(-0.536, 0.655)<br>p = 0.845 |
| Risk Attitude |  | -0.368<br>(-0.799, 0.063)<br>p = 0.094 | -0.143<br>(-0.580, 0.294)<br>p = 0.521 | -0.367<br>(-0.799, 0.065)<br>p = 0.095 | -0.133<br>(-0.572, 0.305)<br>p = 0.550 |
| OT X Disposition to trust |  |  |  | 0.103<br>(-0.758, 0.965)<br>p = 0.813 | 0.280<br>(-0.553, 1.114)<br>p = 0.509 |
| Constant | 7.196<br>(6.585, 7.806)<br>p < 0.001 | 7.174<br>(6.566, 7.782)<br>p < 0.001 | 7.272<br>(6.689, 7.855)<br>p < 0.001 | 7.176<br>(6.567, 7.785)<br>p < 0.001 | 7.280<br>(6.696, 7.864)<br>p < 0.001 |
| **Other personality traits** | **NO** | **NO** | **YES** | **NO** | **YES** |
| R<sup>2</sup> | 0.017 | 0.032 | 0.135 | 0.032 | 0.136 |
| F Statistic | 6.22*<br>(df = 1, 357) | 3.92**<br>(df = 3, 355) | 5.41***<br>(df = 10, 348) | 2.95*<br>(df = 4, 354) | 4.95*** (df = 11, 347) |

\* $p < 0.05$ , \*\* $p < 0.01$ , \*\*\* $p < 0.001$

The results of these regressions show that OT has a robust and significant causal impact on trusting behavior regardless of the covariates that we add. Interestingly, variations in dispositional trust and risk attitudes are not significantly associated with the amount invested in our subject pool of low-trusting individuals (model 1). Perhaps the very fact of being a low-trusting individual “swamps” any variation in dispositions to trust and take risks.

Of particular interest is the question whether there is an interaction effect between OT and individuals’ disposition to trust. One might speculate, for example, that the impact of OT becomes stronger at lower dispositional trust levels. To test this, we illustrate the extent to which OT has a different impact across different levels of the disposition to trust in Figure 2. It documents that a higher disposition to trust is generally correlated with higher behavioral trust levels but the difference between the OT and the placebo condition is nearly constant across individuals’ disposition to trust. For example, Figure 2a shows that, for the lowest disposition to trust (i.e., for dispositional trust scores between 12-23), individuals who received OT transferred roughly €0.7 more to the trustee while for the highest disposition to trust (i.e., for dispositional trust scores between 34-38), the individuals transfer €0.81 more if they received OT instead of placebo. We further validate the regularities suggested by Figure 2a by including an interaction term (OT X disposition to trust) in models 3 and 4 of Table 1. These interaction terms are small and positive in both regressions (0.103 and 0280, respectively) but clearly insignificant (p = 0.813 and p = 0.509, respectively).

**Figure 2.**
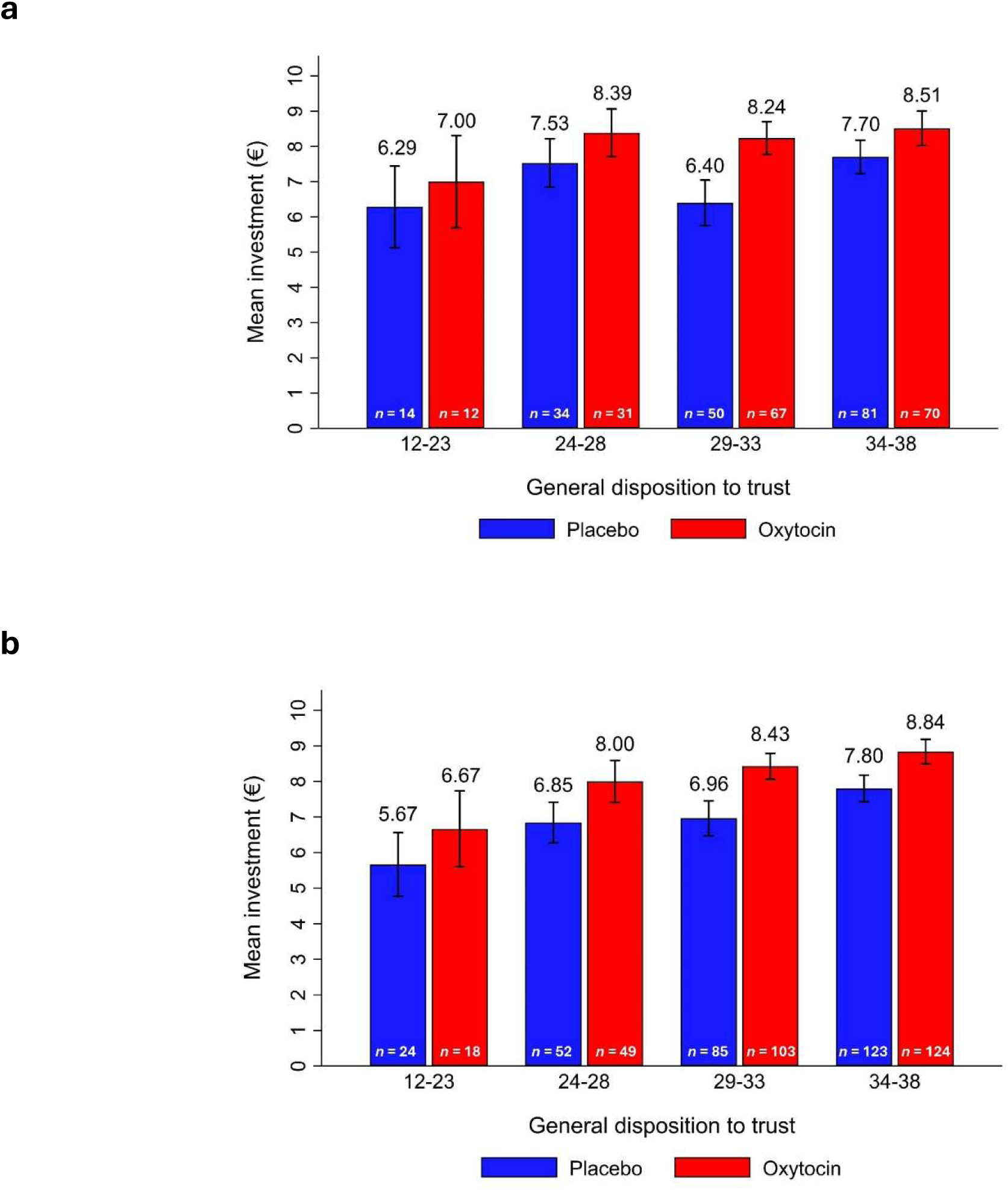
Mean investment (€) by treatment condition and general disposition to trust. Participants were grouped by their general disposition to trust in intervals of 5 points, except for the first group (12-23) which was expanded due to the low number of cases in the lower range of the scale. Mean investments are reported above each bar and sample sizes (n) are shown for each bar. Error bars represent the standard errors of the means in the respective dispositional trust range. **a.** Data from Subjects of the current study (n = 359). **b.** Pooled sample (n = 578) comprising the individuals from our new study and the low-trusting individuals of the no-contract condition in Declerck et al.^8^. For both samples the graphs indicate that at every interval of dispositional trust, the OT group displays a higher level of behavioral trust than the placebo group. In addition, the difference between the OT and the placebo condition is not systematically affected by the disposition to trust.

Turning to the other control variables (see SI, Supplementary Table S1), we note that feelings of connectedness, honesty/humility and, perhaps surprisingly, anxiety have a positive impact on trust while conscientiousness has a negative impact. In contrast, the personality variables extraversion, agreeableness and openness have no significant impact.

We also performed several other robustness checks to make sure that the results cannot be ascribed to mood changes during the course of the experiment, or to participants’ beliefs about whether they had received placebo or OT. (see SI, Supplementary Tables S2 and S3). None of these additional checks questioned the robust influence of OT on subjects’ trusting behavior.

## RESULTS based on the pooled sample (n=578)

The pooled sample merges the data from our new study (n = 359) and the n = 219 individuals with a low trust disposition in the no-contact condition of Declerck et al.^8^. Figure 3 shows the cumulative distributions of dispositional trust scores from these two samples. The figure indicates that the two distributions are rather similar and a two-way ANOVA confirms this impression statistically. There are no significant differences in dispositional trust scores between the oxytocin and placebo conditions (F(1, 574) = 0.28, p = 0.600), between the current experiment and the Declerck et al.^8^ experiment (F(1, 574) = 0.09, p = 0.768) nor do we find an interaction (F(1, 574) = 2.20, p = 0.139).

**Figure 3.**
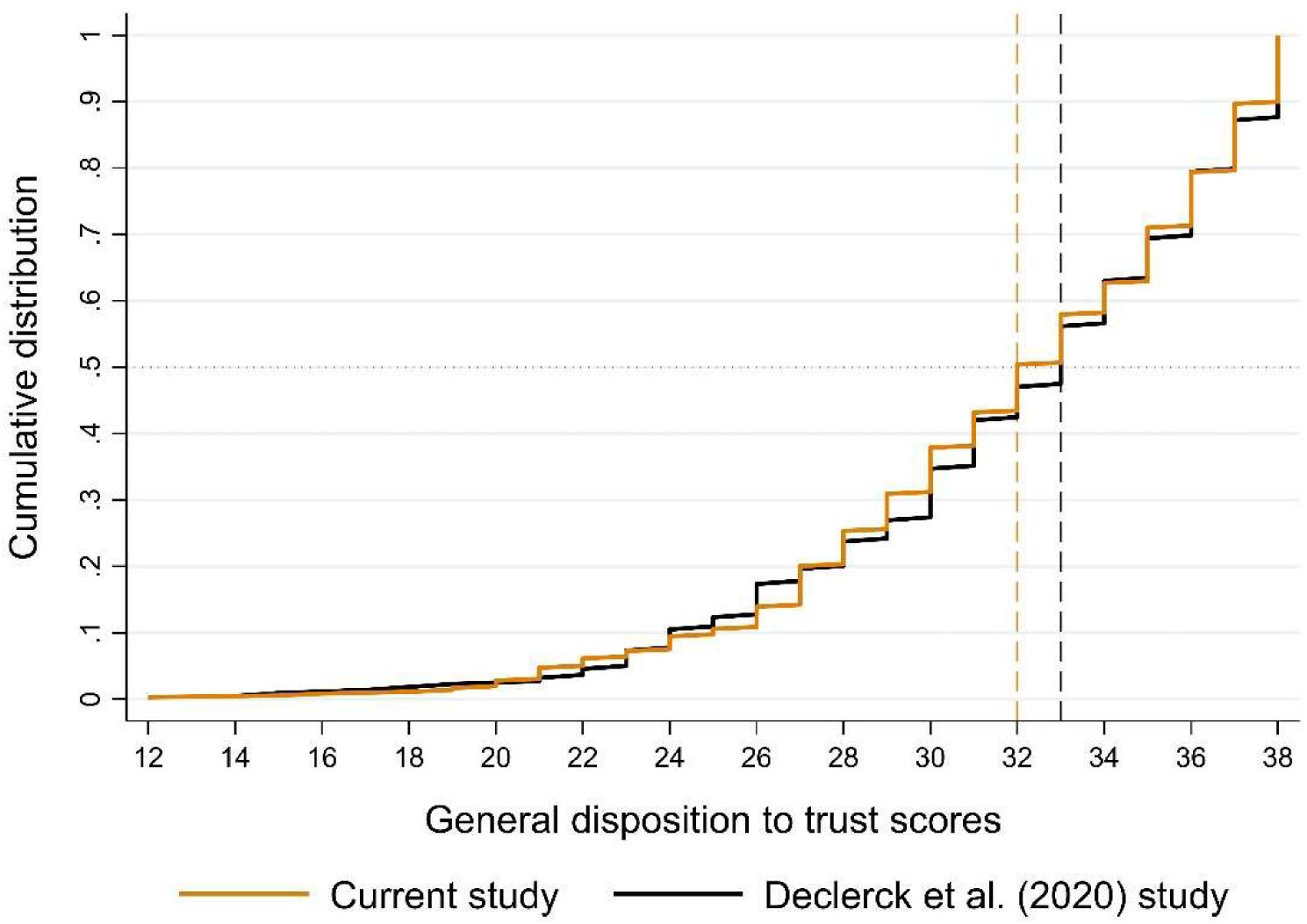
Cumulative distribution of the general disposition to trust scores in the current study and among the low-trusting individuals in the no-contract condition of Declerck et al.^8^. The dashed lines represent the medians of the distributions. The two distributions are very similar and statistically indistinguishable (Mann Whitney U Test, z = 0.475, p = 0.635). The total range of the dispositional trust scale of Yamagishi et al.^25^ is 9-63 while the range of dispositional trust scores for low trusting individuals in the two samples is 12-38.

In the pooled sample, the investors’ mean transfer in the trust experiment is 8.42 (s.d.=3.94) in the OT condition (n=294), while it is only 7.20 (s.d.=4.32) in the placebo condition (n=284), indicating a 16.9% increase in behavioral trust when subjects received OT. This difference is statistically significant at P=0.0004, (two sample t-test, t=-3.56, two-tailed). Figure 2b further strengthens the evidence for a causal effect of OT on trust in low-trusting individuals. The figure documents that at every interval of the dispositional trust score, individuals with OT display a higher level of behavioral trust compared to those with placebo.

The causal role of trust is corroborated by a more detailed statistical analysis that controls for other covariates including subjects’ personality traits (see Table 2 and SI, Supplementary Table S4). In all four models estimated we find a highly significant OT-induced 16 – 17% increase in trusting behaviour. Interestingly, individuals’ disposition to trust is significantly positively correlated with behavioral trust in Model 2 and Model 3 but becomes insignificant in Model 4, where we control for openness, conscientiousness, agreeableness and extraversion. With the exception of conscientiousness (see SI, Supplementary Table S4) – which is negatively correlated with behavioral trust – none of these personality variables is significantly correlated with trust.

**Table 2.** The impact of oxytocin (OT) on investors’ transfers when controlling for covariates (pooled data, n = 578)

|  | Model 0 | Model 1 | Model 2 | Model 3 | Model 4 |
| --- | --- | --- | --- | --- | --- |
| OT | 1.225<br>(0.550, 1.899)<br>p < 0.001 | 1.223<br>(0.555, 1.892)<br>p < 0.001 | 1.173<br>(0.512, 1.833)<br>p = 0.001 | 1.223<br>(0.554, 1.892)<br>p < 0.001 | 1.171<br>(0.510, 1.832)<br>p = 0.001 |
| Disposition to trust |  | 0.597<br>(0.262, 0.931)<br>p < 0.001 | 0.475<br>(0.140, 0.810)<br>p = 0.006 | 0.531<br>(0.063, 0.999)<br>p = 0.026 | 0.382<br>(-0.083, 0.847)<br>p = 0.107 |
| Risk Attitude |  | -0.118<br>(-0.453, 0.217)<br>p = 0.489 | -0.119<br>(-0.451, 0.215)<br>p = 0.486 | -0.118<br>(-0.453, 0.217)<br>p = 0.490 | -0.117<br>(-0.450, 0.217)<br>p = 0.491 |
| OT X Disposition to trust |  |  |  | 0.134<br>(-0.536, 0.803)<br>p = 0.695 | 0.189<br>(-0.471, 0.850)<br>p = 0.574 |
| Constant | 7.197<br>(6.717, 7.678)<br>p < 0.001 | 7.198<br>(6.721, 7.675)<br>p < 0.001 | 7.224<br>(6.754, 7.693)<br>p < 0.001 | 7.197<br>(6.720, 7.674)<br>p < 0.001 | 7.224<br>(6.754, 7.693)<br>p < 0.001 |
| **Other personality traits** | **NO** | **NO** | **YES** | **NO** | **YES** |
| R<sup>2</sup> | 0.022 | 0.043 | 0.085 | 0.044 | 0.086 |
| F Statistic | 12.72***<br>(df = 1, 576) | 8.65***<br>(df = 3, 574) | 6.61***<br>(df = 8, 569) | 6.52***<br>(df = 4, 573) | 5.90***<br>(df = 9, 568) |

\* $p < 0.05$ , \*\* $p < 0.01$ , \*\*\* $p < 0.001$

Is there an interaction effect between OT and the disposition to trust in the pooled sample? Table 2 and Figure 2b indicate that no interaction effect exists. For example, for the lowest disposition to trust (i.e., dispositional trust scores between 12-23) individuals who received OT transferred roughly €1 more to the trustee while for the highest disposition to trust (i.e., dispositional trust scores between 34-38), the individuals in our pooled sample transfer €1.04 more if they received OT instead of a placebo. The absence of an interaction effect is further supported by the insignificant coefficient of the interaction term (OT X disposition to trust) in Models 2-4 of Table 2. Although positive, the interaction term is clearly insignificant in all regressions.

Finally, we performed a similar analysis as in Tables 1, 2 and Figure 2, but separately for the low trusting individuals in the no-contact condition of Declerck et al.^8^ (see SI, Supplementary Table S5 and Figure S1). These additional analyses again corroborate our main finding in the current study: *among low-trusting individuals* those with very low dispositions to trust respond equally strong to OT than those with the highest disposition to trust.

In view of this finding, one might speculate how OT affects the behavioral trust of individuals with a high disposition to trust *when there is no ceiling effect*. If the impact of OT on behavioral trust is not diminished for individuals with an above median disposition to trust, then it should be possible to identify this effect with an experiment that credibly removes the ceiling effect.

## Conclusion

This study provides robust evidence that intranasal oxytocin increases trusting behavior in individuals with a low disposition to trust. By employing a preregistered, high-powered experimental design and targeting a well-defined population, we were able to overcome many of the methodological limitations that have plagued previous oxytocin research. Our findings show a consistent and statistically significant increase in trust among low-trusting individuals following OT administration—an effect replicated in both the standalone sample and a pooled analysis incorporating earlier data.

Importantly, we find no interaction between OT and levels of dispositional trust within the low-trust group, suggesting the effect is relatively uniform across this subpopulation. This pattern contrasts with the ceiling effects seen in high-trusting individuals and underscores the need for future research to examine trust-enhancing interventions in context-specific and population-specific ways.

Our results strengthen the case for a more nuanced understanding of oxytocin’s social effects. Rather than acting as a universal trust enhancer, OT appears to amplify trust where baseline levels are low—much like an analgesic that is only effective in the presence of pain. Future research should explore the neurobiological and psychological mechanisms underlying this selective responsiveness, and test whether similar effects occur in other contexts where social risk and trust are critical. In addition, future research may benefit from further dissecting individual differences that moderate OT’s effects, such as attachment styles, early life experiences, or genetic markers.

## Methods

This Study was approved by the Medical Ethics Commissions of the Otto von Guericke University Magdeburg.

### Participants

A total of 359 male participants took part in this double-blind, placebo-controlled study. The sample size calculation is based on the effect size observed in Declerck et al.^8^ (see Supplementary Table 5) which reflects the effect of oxytocin on investments in the trust game among participants with a total score of 38 on the General Disposition to Trust scale developed by Yamagishi et al. (2015). With this effect size, a minimum of 358 participants (i.e., 179 per condition) is required to achieve a power of 0.95 with a one-sided significance level of 0.05 using a two-sample t-test (See Supplementary information, Table S6).

Participants were students from different faculties at Otto von Guericke University Magdeburg and Leipzig University in Germany. They were initially recruited by distributing links to a paid online questionnaire in lecture halls across the two universities. The online questionnaire was conducted approximately one week before the laboratory experiment where subjects participated in the trust game. It primarily served to identify individuals with low disposition to trust using the General Disposition to Trust scale^25^. Furthermore, during the online questionnaire we also measured general risk attitudes^27^, the Honesty-Humility dimension of the HEXACO scale^26^, and the Anxiety subdimension of the HEXACO scale which measures the general tendency to worry about things. Note that generally worrying about things is not the same as the feeling that one cannot, in general, trust other people because the correlation between individuals’ dispositional trust and their anxiety (as defined in the Hexaco scale) is insignificant and close to zero in our sample. Only participants scoring 38 or lower on the dispositional trust scale were invited to the laboratory experiment. The experiment was advertised as investigating “the psychobiological foundations of decision making” and it took place from November 2023 until January 2025. The study was limited to male participants to be as close as possible to the study design of Declerck et al.^8^ and Kosfeld et al.^1^.

### Experimental procedure

Upon arriving at the laboratory, participants were randomly assigned to either the oxytocin or the placebo condition in a double-blind manner and individually seated in cubicles to prevent visual contact with potential interaction partners. Participants were assigned a cubicle number that served as their personal identification code throughout the experiment. After being seated, participants were asked whether they meet any of the following exclusion criteria: (1) current diagnosis of psychiatric disorders, (2) recurrent problems with substance abuse, (3) current use of psychoactive medications (sleep medication, anxiolytics, antipsychotics, or antidepressants), (4) cold symptoms or severe nasal obstruction, (5) consumption of more than 1 liter of fluid within one hour prior to the experiment, or (6) non-compliance with pre-study abstinence requirements (i.e., no alcohol, non-prescription drugs, or heavy smoking >20 cigarettes for at least 12 hours). Participants were informed that if they were unable to participate due to meeting any exclusion criteria or for any other reason, they would still receive a show-up fee as compensation. After confirming eligibility, all participants read and signed an informed consent form. Following this, participants completed a mood assessment questionnaire (MDMQ)^28^.

According to their random assignment, participants then received either a nasal metered spray of 1 ml syntocinon (with 24 IU of synthetic oxytocin) or placebo (isotonic solution with no active ingredient) each one of them administered with three spray puffs into each nostril. Based on the guidelines of Guastella et al.^29^, detailed oral and written instructions on how to use the spray were given so that the substance would reach the posterior upper end of the nasal cavity (see Supplementary Information Table S7). A supervisor watched as participants self-administered the substance, making sure the instructions were correctly followed. Next participants waited **50 min** after first puff for the active ingredient to take effect^14^. During the waiting period participants first filled out a questionnaire that was based on the 60-items HEXACO questionnaire^26^ and Raven’s task as a filler task. A second mood (MDMQ) and arousal questionnaire was administered approximately 30 minutes after the first puff. After approximately 35 minutes, participants received written instructions for the trust game^30^, which were identical to the instructions in Declerck et al.^8^. Participants were given time to thoroughly read the instructions and complete comprehension questions. Supervisors checked participants’ answers and provided explanations if needed. Participants were told that they would be randomly matched to someone sitting in the other room but that neither their own, nor the partner’s identity would ever be revealed (see Supplementary Methods for the instructions).

The entire laboratory experiment, including the administration of scales and the trust game, was programmed in oTree and conducted on computers linked in a local network. The game began approximately 50 minutes after the first puff. In the trust game, both the investor and trustee received an initial endowment of 12 Euros. The investor had the option to send 0, 4, 8, or 12 Euros to the trustee. Then the experimenter tripled each euro transferred by the investor, adding the trippled amount to the trustee’s initial 12 Euro endowment. Subsequently, the trustee had the option to send back any amount (which was not tripled) ranging from zero to their total available sum.

Each participant played the game twice, first as an investor and then as a trustee, each time with a different partner. The first game in the role of investor occurred without knowledge that there would be a second game in the role of a trustee, and no feedback was given between games. The second game served the purpose to generate trustee decisions that were used to calculate the final payoff of the trustors.

### Statistical analysis

All statistical analyses were conducted using Stata 17.0. The primary hypothesis was tested using a one-tailed two-sample t-test comparing mean investments between the oxytocin and placebo groups, with significance level set at α = 0.05. To provide additional robustness, we conducted supplementary OLS regression analyses using a hierarchical approach with five models (see Table 1), with trust game investments as the dependent variable and treatment condition (Oxytocin vs. placebo) as the primary independent variable. All continuous control variables were z-scored (*M* = 0, *SD* = 1). For the regression analyses, we used two-tailed tests with α = 0.05. To examine the overall robustness of our findings and to increase statistical power, we conducted additional analyses combining data from the current study (n = 359) with the data from participants (n = 219) who had low dispositional trust scores (≤ 38) in Declerck et al.^8^.

## Data availability

All data collected for this study, along with the experimental logs and protocols will be deposited at the Open Science Framework and can then be accessed at osf.io/cu9dx.

## Code availability

The experimental code (oTree implementation) and analysis code will also be made available at osf.io/cu9dx.

## Acknowledgements

The authors did not receive specific funding for this project.

## Author Contributions

C.D., E. F. and B. V. conceptualized and designed the study, P. B. and B. V. conducted the experiment, P. B. analyzed the data with input from C. D., E. F. and B. V.. C. D., E. F., and B. V. wrote the paper with input from P. B.

## Supplementary Information

**Supplementary Table 1.** Linear regression analyses testing for the impact of oxytocin (OT) on investors’ transfers in the current study, with controls for covariates (n = 359)

|  | Model 0 | Model 1 | Model 2 | Model 3 | Model 4 |
| --- | --- | --- | --- | --- | --- |
| OT | 1.093<br>(0.231, 1.955)<br>p = 0.013 | 1.136<br>(0.277, 1.995)<br>p = 0.010 | 0.942<br>(0.114, 1.769)<br>p = 0.026 | 1.136<br>(0.276, 1.996)<br>p = 0.010 | 0.936<br>(0.109, 1.764)<br>p = 0.027 |
| Disposition to trust |  | 0.328<br>(-0.104, 0.759)<br>p = 0.136 | 0.199<br>(-0.225, 0.624)<br>p = 0.357 | 0.277<br>(-0.328, 0.882)<br>p = 0.369 | 0.059<br>(-0.536, 0.655)<br>p = 0.845 |
| Risk Attitude |  | -0.368<br>(-0.799, 0.063)<br>p = 0.094 | -0.143<br>(-0.580, 0.294)<br>p = 0.521 | -0.367<br>(-0.799, 0.065)<br>p = 0.095 | -0.133<br>(-0.572, 0.305)<br>p = 0.550 |
| OT X Disposition to trust |  |  |  | 0.103<br>(-0.758, 0.965)<br>p = 0.813 | 0.280<br>(-0.553, 1.114)<br>p = 0.509 |
| Inclusion of others (IOS) |  |  | 0.662<br>(0.249, 1.075)<br>p = 0.002 |  | 0.657<br>(0.243, 1.071)<br>p = 0.002 |
| Honesty-Humility |  |  | 0.731<br>(0.310, 1.152)<br>p = 0.001 |  | 0.737<br>(0.315, 1.158)<br>p = 0.001 |
| Anxiety |  |  | 0.563<br>(0.141, 0.985)<br>p = 0.009 |  | 0.571<br>(0.148, 0.994)<br>p = 0.008 |
| Openness |  |  | 0.008<br>(-0.410, 0.426)<br>p = 0.968 |  | 0.008<br>(-0.410, 0.427)<br>p = 0.968 |
| Conscientiousness |  |  | -0.515<br>(-0.948, -0.081)<br>p = 0.020 |  | -0.526<br>(-0.961, -0.091)<br>p = 0.018 |
| Agreeableness |  |  | -0.183<br>(-0.604, 0.238)<br>p = 0.393 |  | -0.173<br>(-0.595, 0.249)<br>p = 0.421 |
| Extraversion |  |  | 0.149<br>(-0.295, 0.592)<br>p = 0.510 |  | 0.137<br>(-0.308, 0.583)<br>p = 0.544 |
| Constant | 7.196<br>(6.585, 7.806)<br>p < 0.001 | 7.174<br>(6.566, 7.782)<br>p < 0.001 | 7.272<br>(6.689, 7.855)<br>p < 0.001 | 7.176<br>(6.567, 7.785)<br>p < 0.001 | 7.280<br>(6.696, 7.864)<br>p < 0.001 |
| R<sup>2</sup> | 0.017 | 0.032 | 0.135 | 0.032 | 0.136 |
| F Statistic | 6.22*<br>(df = 1, 357) | 3.92**<br>(df = 3, 355) | 5.41***<br>(df = 10, 348) | 2.95*<br>(df = 4, 354) | 4.95*** (df = 11, 347) |

\* $p < 0.05$ , \*\* $p < 0.01$ , \*\*\* $p < 0.001$

**Supplementary Table 2.** Population-averaged generalized estimating equations (GEE) with exchangeable correlation structure were used to analyze mood changes within experimental sessions. The sample comprised 718 observations from 359 participants. Robust 95% confidence intervals for regression coefficients are shown in parentheses. The 30 items of the mood scale (which were administered before and after oxytocin administration) were categorized in 3 dimensions according to valence (good/bad, 10 items), alertness (awake/tired, 10 items), and calmness (calm/nervous, 10 items). The Good/Bad dimension (first dependent variable) consists of the following items of the MDMS questionnaire 3 (r = reversed item): 1 (content), 4r (bad), 8 (great), 11r (uncomfortable), 14 (superb), 17 (good), 19r (unhappy), 21 (discontent), 24 (happy), 29 (wonderful). The Awake/Tired dimension (second dependent variable) consists of the following items: 2 (rested), 5r (worn out), 7r (tired), 10 (energetic), 13 (activated), 16r (sleepy), 20 (alert), 23 (fresh), 26r (exhausted), 28 (wide awake). The Calm/Nervous dimension (third dependent variable) consists of the following items: 3r (restless), 6 (composed), 9r (uneasy), 12 (relaxed), 15 (absolutely calm), 18 (at ease), 22r (tense), 25r (nervous), 27 (calm), 30 (deeply relaxed). The variable Time takes on the value 0 if the items were assessed prior to the time at which oxytocin administration took place, and 1 indicates the mood assessment that took place after the time of OT administration.

|  | **Good/Bad** | **Awake/Tired** | **Calm/Nervous** |
| --- | --- | --- | --- |
| Oxytocin | -0.076<br>(-0.207, 0.055)<br>*p* = 0.253 | -0.066<br>(-0.226, 0.093)<br>*p* = 0.416 | -0.101<br>(-0.246, 0.044)<br>*p* = 0.170 |
| Time | -0.059<br>(-0.126, 0.007)<br>*p* = 0.081 | -0.099<br>(-0.176, -0.023)<br>*p* = 0.011 | 0.079<br>(-0.002, 0.161)<br>*p* = 0.056 |
| Oxytocin x Time | 0.026<br>(-0.060, 0.113)<br>*p* = 0.548 | -0.051<br>(-0.176, -0.023)<br>*p* = 0.376 | 0.019<br>(-0.092, 0.130)<br>*p* = 0.738 |
| Constant | 4.650<br>(4.553, 4.746)<br>*p* < 0.001 | 4.310<br>(4.193, 4.427)<br>*p* < 0.001 | 4.440<br>(4.332, 4.548)<br>*p* < 0.001 |
| Observations | 359 x 2 | 359 x 2 | 359 x 2 |
| *Pseudo-R*<sup>2</sup> | 0.004 | 0.009 | 0.008 |

The results indicate that mood changes during the course of the experiment were primarily time-dependent rather than influenced by oxytocin administration. Participants reported feeling less alert the second time they filled out the questionnaire, as the significant time coefficient of the Awake/Tired dimension (*p* = 0.011) suggests. A trend toward feeling worse over time was also noted in the Good/Bad dimension (*p* = 0.081), as well as a trend in the Calm/Nervous dimension (*p* = 0.056). However, oxytocin itself did not significantly impact any of the three mood dimensions, nor was there a significant interaction between oxytocin and time.

**Supplementary Table 3a.** Participants’ post-treatment beliefs about whether they received oxytocin or placebo. While more participants believed they received placebo, this tendency did not differ significantly between actual treatment groups (Pearson chi²(1) = 0.8123, *p* = 0.367). The remaining 142 participants responded with “I don’t know.”

| Treatment | Beliefs |  | Total (column) |
| --- | --- | --- | --- |
|  | Assumed placebo | Assumed hormone |  |
| Placebo | 75<br>(46.58%) | 30<br>(53.57%) | 105<br>(48.39%) |
| Oxytocin | 86<br>(53.42%) | 26<br>(46.43%) | 112<br>(51.61%) |
| Total (row) | 161<br>(100%) | 56<br>(100%) | 217<br>(100%) |

*Note.* Percentages in the "Assumed placebo" and "Assumed hormone" columns represent column percentages. The remaining 142 participants responded with “I don't know” or left the answer blank.

**Supplementary Table 3b.** Linear regression analysis of the impact of treatment beliefs on investment choices. “Assumed hormone” is a binary variable with 1 if the participant believed he received hormone and 0 if he believed he received placebo or stated “I don’t know.” According to the results, there is no evidence for a placebo effect. 95% confidence intervals of the regression coefficients are reported in parentheses.

|  | Model<br>controlling for<br>beliefs | Model 1 from<br>Table 1 in<br>manuscript |
| --- | --- | --- |
| Oxytocin | 1.139<br>(0.279, 2.000)<br>$p = 0.010$ | 1.136<br>(0.277, 1.995)<br>$p = 0.010$ |
| General disposition to trust | 0.326<br>(-0.106, 0.758)<br>$p = 0.139$ | 0.328<br>(-0.104, 0.759)<br>$p = 0.136$ |
| Risk attitude | -0.368<br>(-0.799, 0.064)<br>$p = 0.095$ | -0.368<br>(-0.799, 0.063)<br>$p = 0.094$ |
| Assumed hormone | 0.143<br>(-1.042, 1.329)<br>$p = 0.944$ | |
| Constant | 7.150<br>(6.510, 7.790)<br>$p = 0.000$ | 7.174<br>(6.566, 7.782)<br>$p = 0.000$ |
| Observations | 359 | 359 |
| $R^2$ | 0.032 | 0.032 |

**Supplementary Table 4.** Linear regression analyses testing the impact of oxytocin (OT) on investors’ transfers controlling for covariates (pooled sample, n = 578).

|  | Model 0 | Model 1 | Model 2 | Model 3 | Model 4 |
| --- | --- | --- | --- | --- | --- |
| OT | 1.225<br>(0.550, 1.899)<br>$p < 0.001$ | 1.223<br>(0.555, 1.892)<br>$p < 0.001$ | 1.173<br>(0.512, 1.833)<br>$p = 0.001$ | 1.223<br>(0.554, 1.892)<br>$p < 0.001$ | 1.171<br>(0.510, 1.832)<br>$p = 0.001$ |
| Disposition to trust | | 0.597<br>(0.262, 0.931)<br>$p < 0.001$ | 0.475<br>(0.140, 0.810)<br>$p = 0.006$ | 0.531<br>(0.063, 0.999)<br>$p = 0.026$ | 0.382<br>(-0.083, 0.847)<br>$p = 0.107$ |
| Risk Attitude | | -0.118<br>(-0.453, 0.217)<br>$p = 0.489$ | -0.119<br>(-0.451, 0.215)<br>$p = 0.486$ | -0.118<br>(-0.453, 0.217)<br>$p = 0.490$ | -0.117<br>(-0.450, 0.217)<br>$p = 0.491$ |
| OT × Disposition to trust | | | | 0.134<br>(-0.536, 0.803)<br>$p = 0.695$ | 0.189<br>(-0.471, 0.850)<br>$p = 0.574$ |
| Inclusion of others (IOS) | | | 0.697<br>(0.367, 1.028)<br>$p < 0.001$ | | 0.7<br>(0.369, 1.030)<br>$p < 0.001$ |
| Openness | | | 0.091<br>(-0.287, 0.485)<br>$p = 0.614$ | | 0.104<br>(-0.282, 0.491)<br>$p = 0.596$ |
| Conscientiousness | | | -0.534<br>(-0.960, -0.112)<br>$p = 0.013$ | | -0.536<br>(-0.958, -0.113)<br>$p = 0.013$ |
| Agreeableness | | | 0.278<br>(-0.106, 0.663)<br>$p = 0.156$ | | 0.281<br>(-0.104, 0.666)<br>$p = 0.153$ |
| Extraversion | | | 0.132<br>(-0.299, 0.562)<br>$p = 0.548$ | | 0.122<br>(-0.310, 0.554)<br>$p = 0.581$ |
| Constant | 7.197<br>(6.716, 7.678)<br>$p < 0.001$ | 7.198<br>(6.721, 7.675)<br>$p < 0.001$ | 7.224<br>(6.754, 7.693)<br>$p < 0.001$ | 7.197<br>(6.720, 7.674)<br>$p < 0.001$ | 7.224<br>(6.754, 7.693)<br>$p < 0.001$ |
| R<sup>2</sup> | 0.022 | 0.043 | 0.085 | 0.044 | 0.086 |
| F Statistic | 12.72***<br>(df = 1, 576) | 8.65***<br>(df = 3, 574) | 6.61***<br>(df = 8, 569) | 6.52***<br>(df = 4, 573) | 5.90***<br>(df = 9, 568) |

\* $p < 0.05$ , \*\* $p < 0.01$ , \*\*\* $p < 0.001$

**Supplementary Table 5.** Linear regression analyses testing the impact of oxytocin (OT) on investors’ transfers controlling for covariates for low-trusting individuals in the no-contact condition of Declerck et al.^2^ (n = 219).

|  | Model 0 | Model 1 | Model 2 | Model 3 | Model 4 |
| --- | --- | --- | --- | --- | --- |
| OT | 1.432<br>(0.339, 2.525)<br>p = 0.010 | 1.236<br>(0.168, 2.305)<br>p = 0.023 | 1.243<br>(0.183, 2.303)<br>p = 0.022 | 1.237<br>(0.166, 2.307)<br>p = 0.024 | 1.245<br>(0.183, 2.307)<br>p = 0.022 |
| Disposition to trust |  | 0.995<br>(0.460, 1.529)<br>p < 0.001 | 0.895<br>(0.363, 1.426)<br>p = 0.001 | 0.926<br>(0.185, 1.668)<br>p = 0.015 | 0.791<br>(0.052, 1.529)<br>p = 0.036 |
| Risk Attitude |  | 0.265<br>(-0.268, 0.798)<br>p = 0.328 | 0.171<br>(-0.358, 0.701)<br>p = 0.525 | 0.263<br>(-0.271, 0.797)<br>p = 0.333 | 0.169<br>(-0.362, 0.700)<br>p = 0.530 |
| OT × Disposition to trust |  |  |  | 0.143<br>(-0.930, 1.215)<br>p = 0.793 | 0.219<br>(-0.856, 1.293)<br>p = 0.689 |
| Inclusion of others (IOS) |  |  | 0.677<br>(0.141, 1.213)<br>p = 0.014 |  | 0.690<br>(0.149, 1.230)<br>p = 0.013 |
| Openness |  |  | 0.149<br>(-0.593, 0.891)<br>p = 0.692 |  | 0.162<br>(-0.584, 0.908)<br>p = 0.669 |
| Conscientiousness |  |  | -0.505<br>(-1.421, 0.411)<br>p = 0.278 |  | -0.492<br>(-1.412, 0.428)<br>p = 0.293 |
| Agreeableness |  |  | 0.841<br>(0.092, 1.590)<br>p = 0.028 |  | 0.829<br>(0.075, 1.582)<br>p = 0.031 |
| Extraversion |  |  | -0.225<br>(-1.096, 0.647)<br>p = 0.612 |  | -0.242<br>(-1.120, 0.635)<br>p = 0.587 |
| Constant | 7.200<br>(6.411, 7.989)<br>p < 0.001 | 7.302<br>(6.533, 8.070)<br>p < 0.001 | 7.298<br>(6.538, 8.058)<br>p < 0.001 | 7.295<br>(6.523, 8.067)<br>p < 0.001 | 7.288<br>(6.525, 8.051)<br>p < 0.001 |
| R<sup>2</sup> | 0.022 | 0.091 | 0.1397 | 0.091 | 0.1404 |
| F Statistic | 12.72***<br>(df = 1, 217) | 7.16***<br>(df = 3, 215) | 4.26***<br>(df = 8, 210) | 5.36***<br>(df = 4, 214) | 3.79***<br>(df = 9, 209) |

\* $p < 0.05$ , \*\* $p < 0.01$ , \*\*\* $p < 0.001$
The dependent variable is the investors' transfer to the trustee in the trust experiment. All continuous independent variables are z-scored with a mean of zero and a SD of 1. Oxytocin (OT) is a binary variable (0 = Placebo, 1 = Oxytocin). *Risk attitude* measures individuals' general propensity to avoid risks. *Disposition to trust* represents individuals' general disposition to trust as measured in Yamagishi et al.<sup>1</sup>. *Inclusion of others (IOS)* measures individuals' feelings of connectedness with their interaction partner in the trust game. In model 4, we also control for openness, conscientiousness, agreeableness and extraversion. Since Declerck et al.<sup>2</sup> did not measure the honesty/humility and the anxiety dimension of the Hexaco scale, we cannot include these variables in this sample. 95% confidence intervals are shown in parentheses.

**Supplementary Table 6.** A priori sample size computation using the G*Power 3.1.9.7 software ^3^

| **t tests - Means: Difference between two independent means (two groups)** |  |  |
| --- | --- | --- |
| **Analysis:** A priori: Compute required sample size |  |  |
| **Input** | Tail(s) | = One |
|  | Effect size d | = 0.3484542 |
| | $\alpha$ err prob | = 0.05 |
| | Power (1-β err prob) | = 0.95 |
|  | Allocation ratio N2/N1 | = 1 |
| **Output:** | Noncentrality parameter $\delta$ | = 3.2965314 |
|  | Critical t | = 1.6491451 |
|  | Df | = 356 |
|  | Sample size group 1 | = 179 |
|  | Sample size group 2 | = 179 |
|  | Total sample size | = **358** |

**Supplementary Table 7.** Guidelines for OT administration, based on recommendations by Guastella et al.^4^ and also used in Declerck et al.^2^.

|  |  |
| --- | --- |
| **1** | *If necessary, clear your nose from any obstruction (box of tissues provided).* |
| **2** | *Prime the bottle and complete a test spray in the air.* |
| **3** | *Sit comfortably and keep the head in an upright position.* |
| **4** | *Close one nostril with one finger while administering the spray to the other nostril.* |
| **5** | *Insert bottle 1 cm into the nostril and keep the tip of the bottle at a 45 degree angle into the nose. Aim towards the upper lateral part of the nose (and not towards the middle of the nose).* |
| **6** | *Upon delivery, inhale and breathe in lightly. Do not sniff exaggeratedly.* |
| **7** | *Alternate administrations between nostrils. Allow time between each re-administration to the same nostril of at least 15 seconds.* |

**Supplementary Figure 1.**
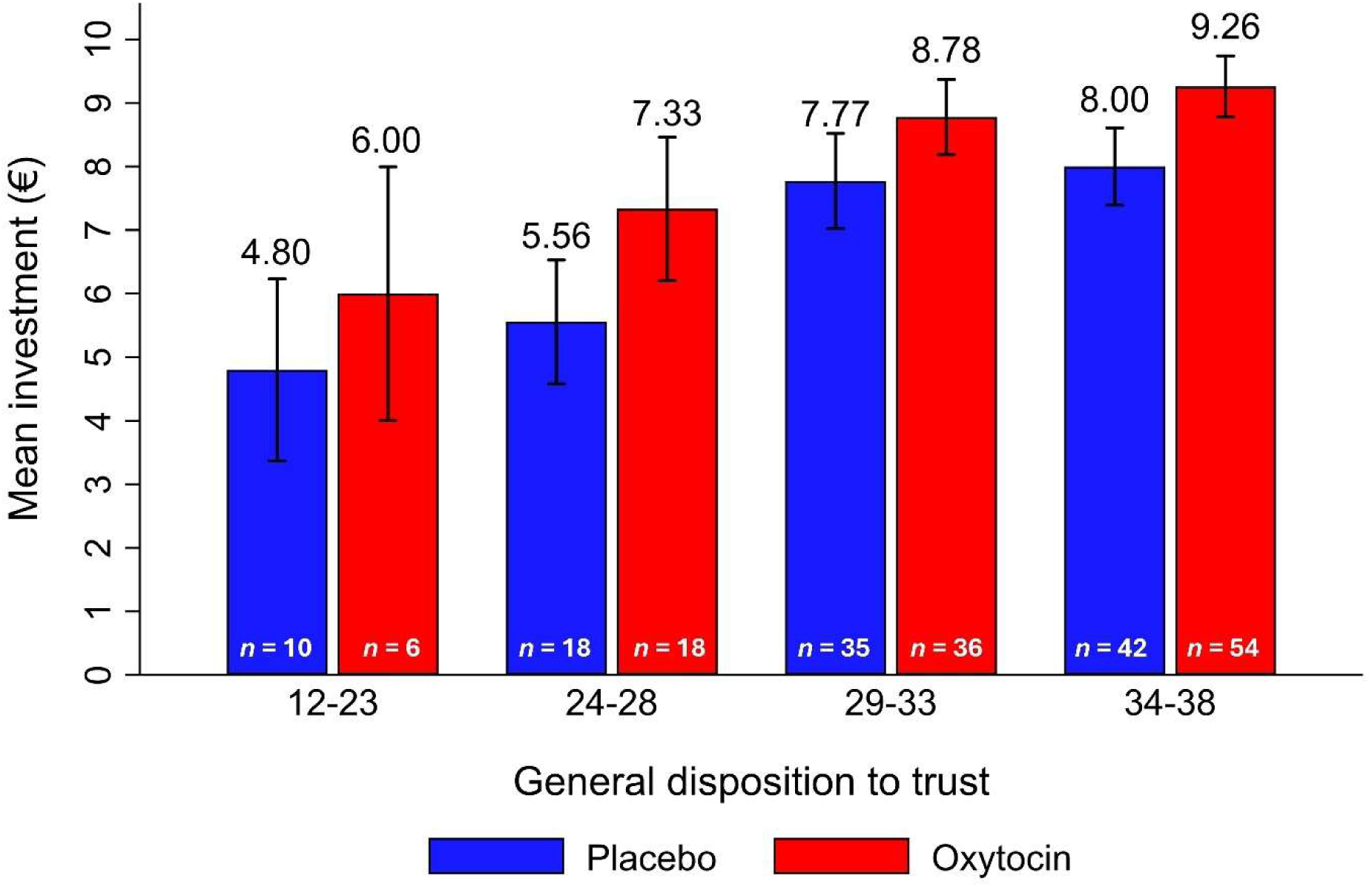
Mean investment (€) by treatment condition and general disposition to trust in the low-trusting population of the no-contact condition (n = 219) of Declerck et al.^2^. Participants are grouped by their general disposition to trust in intervals of 5 points, except for the first group (12-23) which was expanded due to the low number of cases in the lower range of the scale. Mean investments are reported above each bar and sample sizes (n) are shown for each bar. Error bars represent standard errors of the means in the respective dispositional trust range. The graph indicates that at every interval of dispositional trust, the OT group displays a higher level of behavioral trust than the placebo group. In addition, the difference between the OT and the placebo condition is not systematically affected by the disposition to trust.

## Supplementary Methods

### Instructions for the trust game

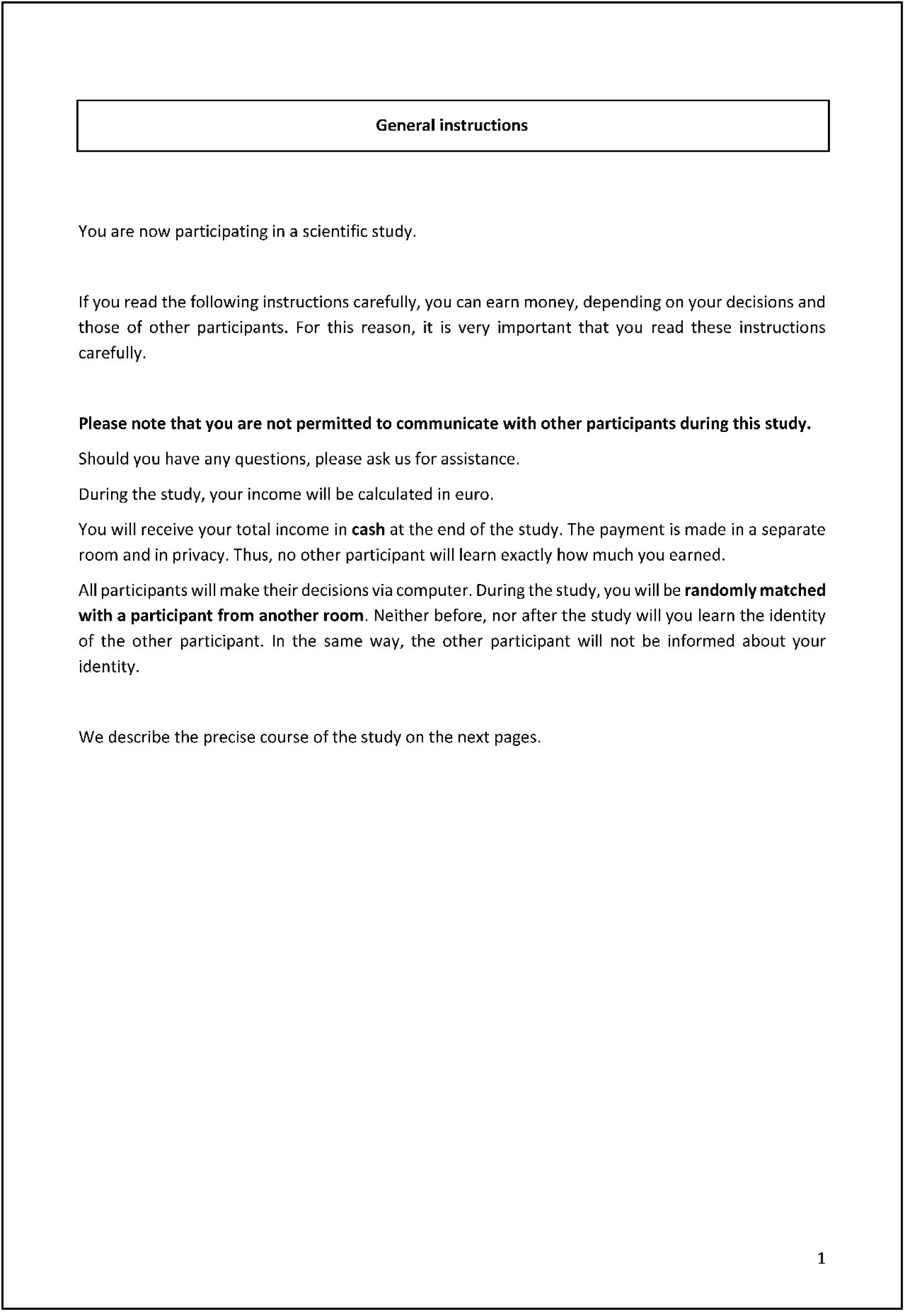
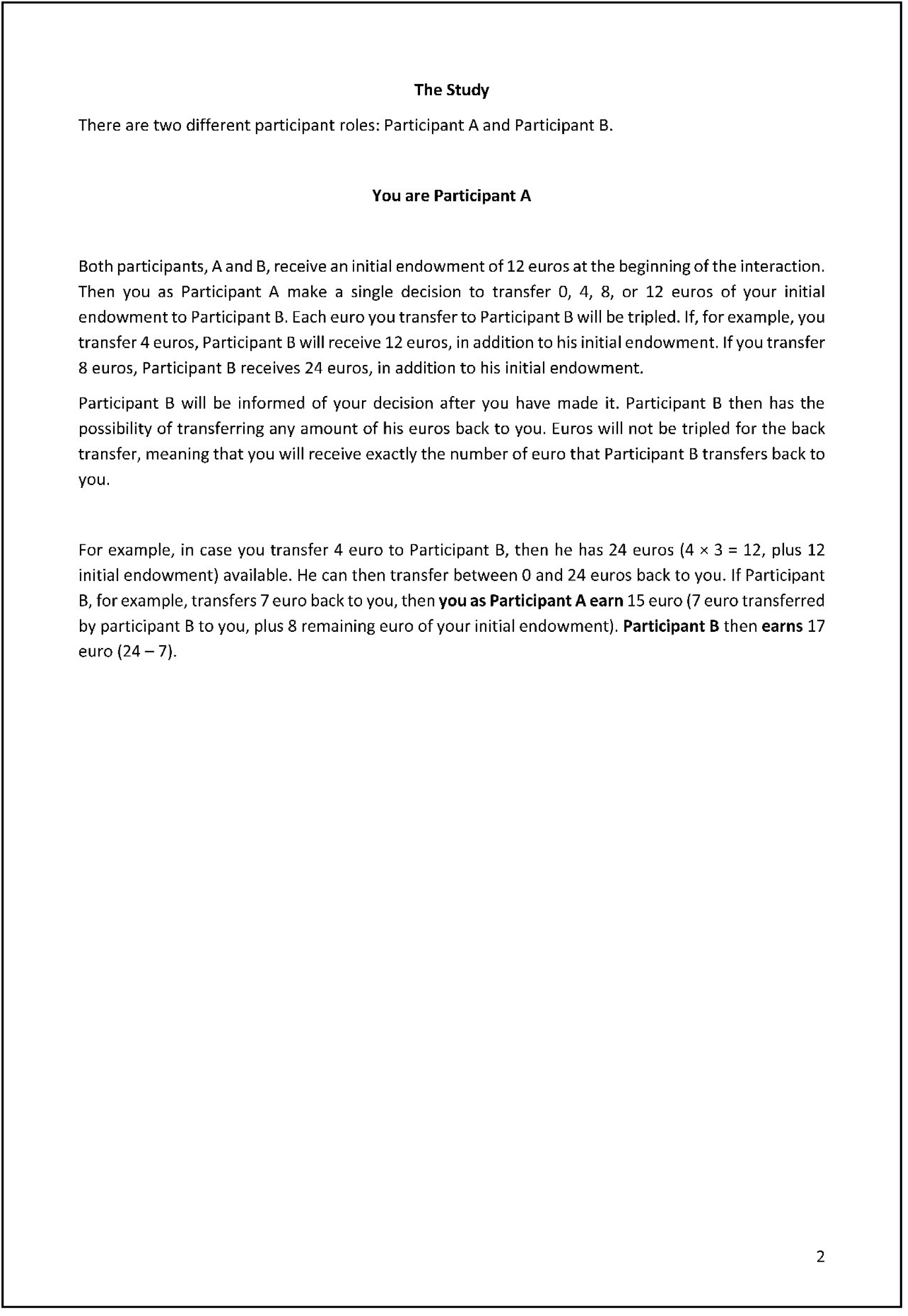
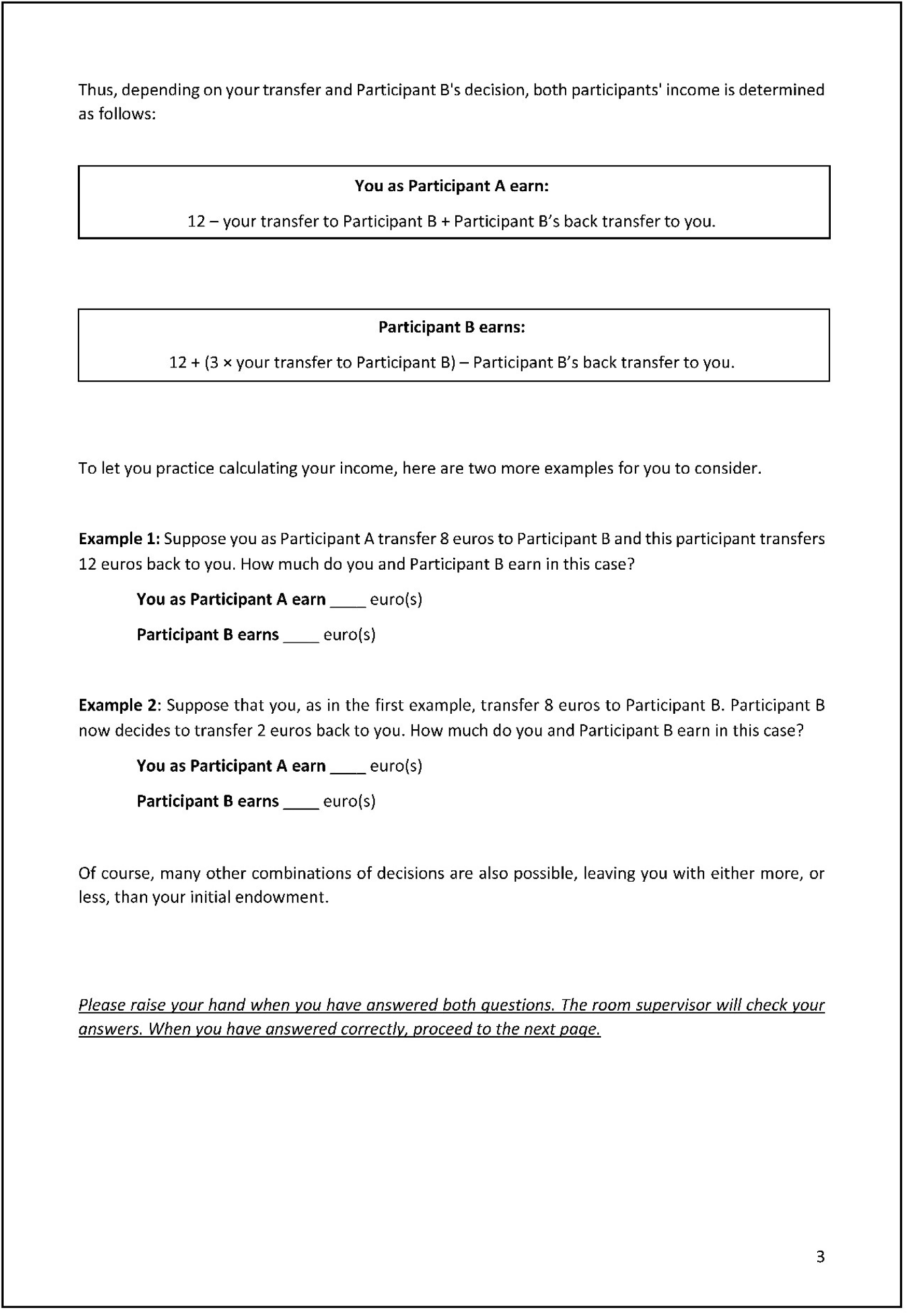
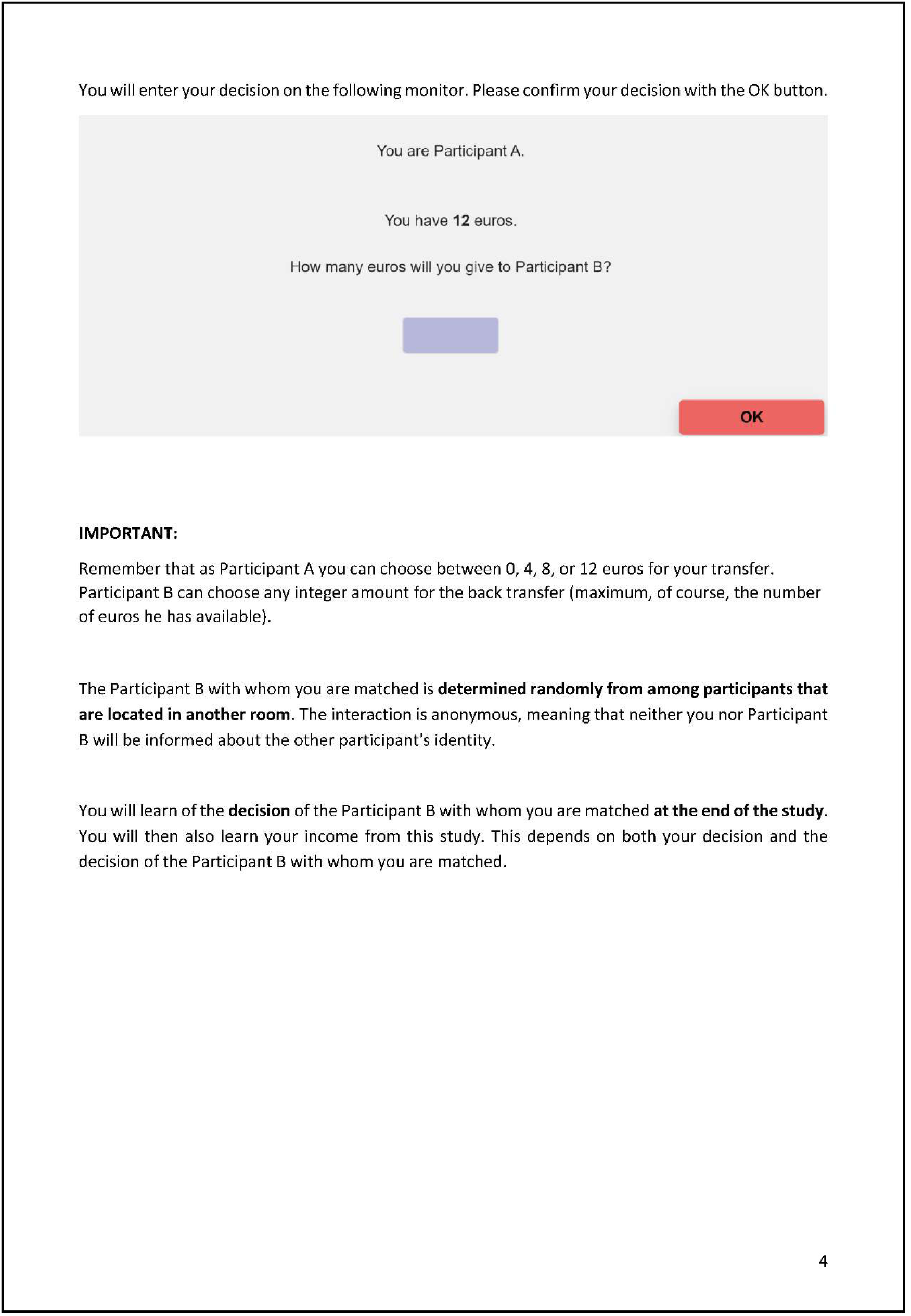

